# Minimally invasive delivery of optical nanosensors using injectable hydrogels

**DOI:** 10.64898/2025.12.21.695747

**Authors:** Shirel Kleiner, Gili Bisker

**Affiliations:** School of Biomedical Engineering, Faculty of Engineering, Tel Aviv University, Tel Aviv 6997801, Israel; Center for Physics and Chemistry of Living Systems, Tel Aviv University, Tel Aviv 6997801, Israel; Center for Nanoscience and Nanotechnology, Tel Aviv University, Tel Aviv 6997801, Israel; Center for Light-Matter Interaction, Tel Aviv University, Tel Aviv 6997801, Israel; Sagol School of Neuroscience, Tel Aviv University, Tel Aviv 6997801, Israel

**Keywords:** single-walled carbon nanotubes, fluorescent sensors, near-infrared, peptide hydrogels, injectable hydrogels

## Abstract

Optimizing treatment plans based on an individual’s drug concentration can significantly improve therapeutic outcomes, as drug efficacy is closely linked to its plasma levels. This motivates the development of precise tools for measuring drug concentration *in situ*. To address this, a minimally invasive optical sensing platform based on preformed, injectable hydrogels that encapsulate near-infrared fluorescent single-walled carbon nanotube (SWCNT) sensors is introduced. DNA-suspended SWCNTs are integrated into self-assembling peptide hydrogels reinforced with natural polysaccharide additives, yielding self-healing, shear-thinning gels that can be injected through a fine needle while maintaining both structural integrity and optical functionality. Using levodopa, the primary treatment for Parkinson’s disease, as a model target analyte, the encapsulated SWCNTs exhibit a dose-dependent fluorescence response across physiologically relevant concentrations. Importantly, the SWCNTs within the hydrogel retain their fluorescence response to levodopa following injection, as well as in the presence of serum, and in a subcutaneous tissue phantom that mimics the mechanical and optical scattering properties of soft tissue. These results establish preformed injectable peptide-based hydrogels as a new class of function-preserving materials for the localized deployment of SWCNT optical nanosensors, enabling minimally invasive *in situ* monitoring of drug levels and offering a potential route toward personalized therapeutic drug monitoring.

## Introduction

The therapeutic efficacy and potential toxicity of drugs are directly linked to their local concentration in the bloodstream or interstitial fluid of various organs and tissues.^[1]^ Monitoring these drug levels enables clinicians to optimize dosing regimens, minimize adverse effects, and improve patient outcomes.^[2]^ This practice, known as therapeutic drug monitoring (TDM), is primarily applied to drugs with narrow therapeutic windows or those that cause severe side effects when under- or overdosed.^[3,4]^ A central challenge in TDM is the development of accurate, reliable, and minimally invasive tools that can measure drug concentrations in real time. Moreover, TDM is typically based on systemic drug concentration measurements, whereas localized, site-specific drug concentrations may better correlate with therapeutic efficacy and clinical outcomes.

Nanosensors offer a promising solution by transducing the interaction between an analyte and a recognition element into a measurable, quantifiable signal.^[5]^ Fluorescent nanosensors, specifically, have attracted considerable attention for continuous physiological monitoring owing to their high sensitivity.^[6,7]^ Among them, single-walled carbon nanotube (SWCNT)-based sensors are particularly attractive for continuous, real-time monitoring, owing to their biocompatibility and unique optical properties.^[8–11]^ Semiconducting SWCNTs fluoresce in the near-infrared (NIR) range, which coincides with the biological transparency window and reduces autofluorescence interference.^[12–18]^ Moreover, their photostability and lack of photobleaching make them suitable for prolonged monitoring.^[19]^ Their surface chemistry and corona phase can be tuned by functionalization with surfactants,^[20]^ DNA,^[21–25]^ peptides,^[26,27]^ or polymers,^[28–30]^ as well as by introducing defects,^[31–40]^ thereby enabling selective interactions with different target molecules that induce fluorescence intensity modulations or wavelength shifts.^[41,42]^

Delivering SWCNT sensors into tissues or blood vessels has been explored through various strategies.^[43,44]^ Direct insertion of SWCNTs was achieved by intravenous injection, which primarily leads to accumulation in the Kupffer cells in the liver,^[45,46]^ by direct injection into tissues such as the liver *ex vivo*^[47]^ and the hippocampus *in vivo*,^[48]^ and by utilizing a reverse electrodialysis battery to deliver functionalized SWCNTs into epidermal and dermal layers of the skin.^[49]^ While straightforward, these methods are limited to the regions of accumulation and carry the risk of off-target uncontrolled diffusion. Alternative strategies rely on surgically implanted SWCNT-containing devices. For example, SWCNTs were either encapsulated as liquid suspensions within a semipermeable dialysis bag^[50,51]^ or a 3D-printed hydrogel core,^[52]^ or integrated within the three-dimensional network of a hydrogel.^[53–58]^ To minimize the invasiveness of the procedure, injectable hydrogels have emerged as a promising platform for *in vivo* delivery of SWCNTs. For instance, preformed hydrogels have been implanted into marine organisms using a large-diameter trocar,^[59]^ and polymeric solutions have been injected into a murine model, forming gels *in situ* following injection within 15 minutes.^[60]^ Despite these significant advances, a delivery system for SWCNT-based sensors that is simultaneously minimally invasive, compatible with administration via a narrow needle, and immediately functional upon *in situ* placement is still needed. Addressing this challenge is thus essential for enabling truly practical, localized, and real-time optical sensing in biological tissues.

Low-molecular-weight peptide gelators are particularly appealing hydrogel platforms due to their biocompatibility, chemical versatility, and low cost.^[61]^ They are characterized by their ability to self-assemble into supramolecular self-supporting hydrogels in response to external stimuli such as solvent, temperature, and pH.^[62,63]^ The three-dimensional network of the hydrogel is formed by physical rather than chemical interactions, namely, π-π stacking, van der Waals forces, and hydrogen bonds. Fluorenylmethoxycarbonyl-diphenylalanine (FmocFF) is one of the most widely studied peptide gelators due to its simplicity, versatility, and ability to self-assemble into an ordered tubular network.^[64,65]^ Hybrid systems incorporating FmocFF with additional components further enhance the mechanical and functional properties of these hydrogels, enabling the tailoring of structural and functional properties for a variety of applications.^[66]^ Therefore, an injectable hydrogel based on self-assembling peptides could offer a versatile injectable platform, as the gelation is reversible and non-covalent.^[67,68]^

Here, we develop a minimally invasive and localized delivery platform of injectable preformed hydrogels encapsulating SWCNTs-based sensors. We utilize single-strand DNA (ssDNA) suspended SWCNTs that show dose-dependent fluorescence response to levodopa (L-DOPA), the primary treatment for Parkinson’s disease (PD), chosen as a model analyte, with a physiologically relevant limit of detection (LOD) below 1 µM. Dosing of L-DOPA is particularly challenging as excessive levels can cause dyskinesia, nausea, and dizziness, whereas insufficient levels reduce therapeutic efficacy. Moreover, as PD progresses, the therapeutic window for L-DOPA narrows, making precise dosing increasingly critical,^[69–72]^ which gives rise to the need for L-DOPA TDM.^[73]^ We integrate the DNA-SWCNTs within a self-assembling FmocFF hydrogel supplemented with either sodium alginate (SA) or hyaluronic acid (HA) to improve injectability. We demonstrate that the resulting hydrogels exhibit shear-thinning and self-healing properties, enabling the fully formed hydrogels to be injected through a fine needle, immediately reassemble while maintaining their structural integrity and the functionality of the embedded nanosensors post-injection. To further establish the utility of the platform, we injected DNA-SWCNT-containing hydrogels into a subcutaneous tissue-mimicking phantom with relevant optical scattering properties and demonstrated the fluorescence response of the encapsulated SWCNT sensors to L-DOPA. Together, the results demonstrate the potential of peptide self-assembling hydrogels as a versatile, minimally invasive *in situ* delivery vehicle for SWCNT-based optical sensing, facilitating the local, real-time monitoring of clinically important analytes utilizing the SWCNT NIR fluorescence.

## Results and discussion

To develop an injectable hydrogel platform for SWCNT-based sensors, we first established a proof-of-concept model system by identifying a functionalized SWCNT formulation with a robust optical response to a small-molecule drug, and then integrating it into self-assembling peptide hydrogels with suitable mechanical properties. SWCNTs suspended with ssDNA are known to exhibit fluorescence modulation in response to small molecules.^[74–76]^ Leveraging this capability, we generated a library of seven ssDNA-wrapped SWCNTs (ssDNA-SWCNTs) (Table S1) and screened their fluorescence responses to our model analyte, the small-molecule drug L-DOPA, which is the precursor of dopamine that is primarily used in the treatment of Parkinson’s disease.^[77]^ Distinct absorption peaks corresponding to various SWCNT chiralities, along with well-defined fluorescence emission peaks in the excitation-emission map, confirmed successful formation of the colloidal suspensions (Figure S1).

Among the tested sequences, the (GT)_15_GATCTAAGGCGTGTAT-SWCNTs (DNA1-SWCNTs) exhibited the most pronounced fluorescence enhancement upon exposure to 1 mM of L-DOPA at a ssDNA-SWCNT concentration of 3 mg mL^-1^ (Figure 1A), and it was therefore chosen as a representative sensor-analyte model system. The (9,4) chirality of the DNA1-SWCNTs, excited at 730 nm and emitting at 1130 nm, showed the highest intensity increase and was selected for further characterization (Figure 1B).

**Figure 1.**
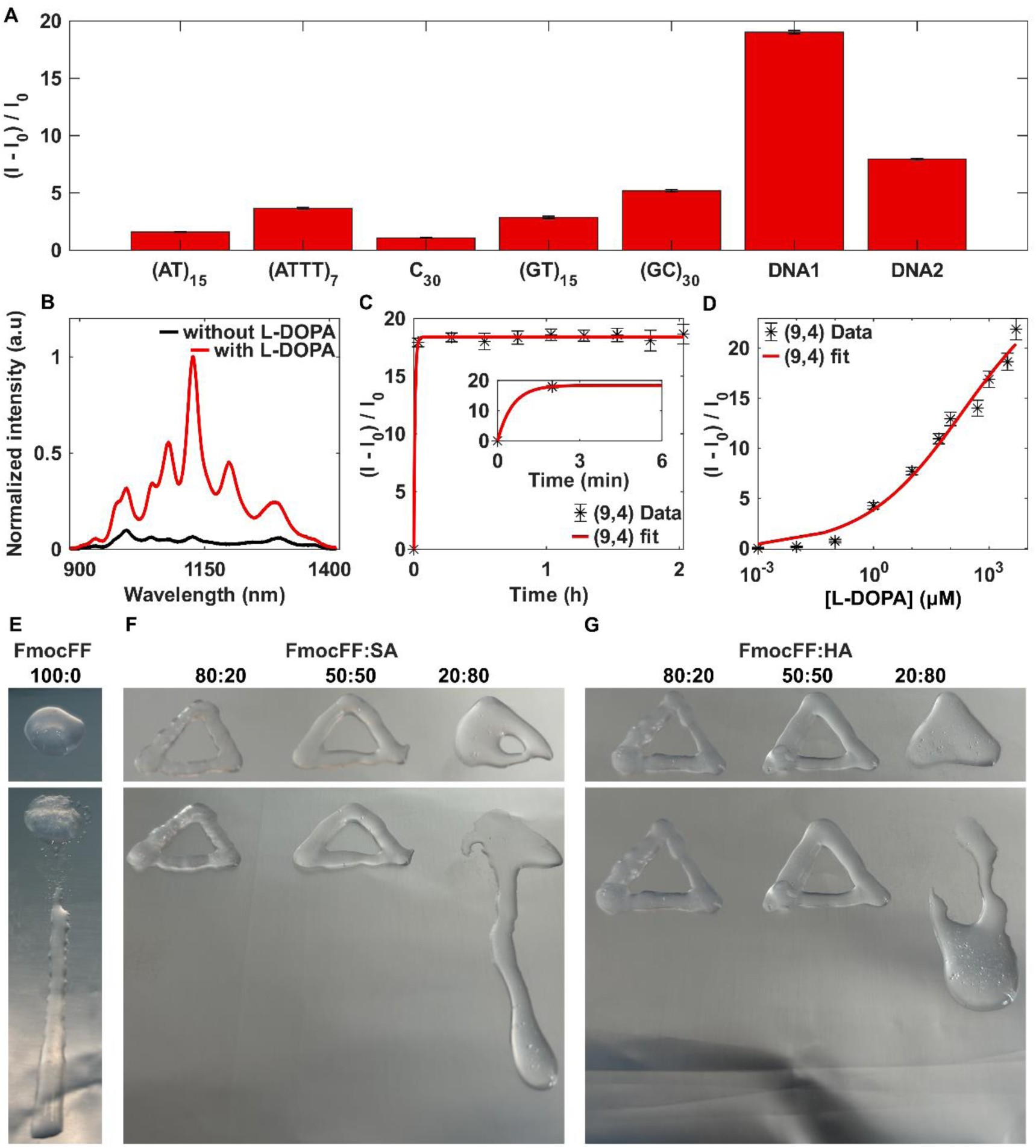
Solution phase characterization and hydrogel injectability. (A) Normalized fluorescence response of ssDNA-SWCNTs library to 1 mM of L-DOPA. (B) Normalized intensity of (GT)_15_GATCTAAGGCGTGTAT-SWCNTs before (black line) and after (red line) the addition of L-DOPA. (C) Time-dependent fluorescence intensity response of the (9,4) chirality of (GT)_15_GATCTAAGGCGTGTAT-SWCNTs to L-DOPA, fitted to a one-phase association equation (Equation 1, red line). Inset: Zoom in to the first six minutes. (D) Dose-dependent fluorescence response of the (9,4) chirality of the DNA1-SWCNTs to increasing concentrations of L-DOPA, fitted by the Hill equation (Equation 2, red line). (E) Injectability of FmocFF hydrogel, (F) Injectability of FmocFF/SA hydrogels with FmocFF:SA ratios of 80:20, 50:50, 20:80, and (G) Injectability of FmocFF/HA hydrogels with FmocFF:HA ratios of 80:20, 50:50, 20:80. Top row: hydrogels immediately after injection. Bottom row: hydrogels after they were tilted to a 90° angle for 10 seconds.

The time-dependent fluorescence of DNA1-SWCNTs upon L-DOPA addition exhibited a rapid response that remained stable over time (Figure 1C). The data were fitted by a one-phase association model:

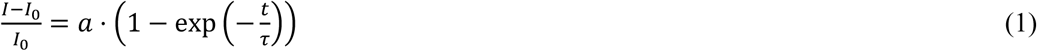

where *I_0_* is the initial fluorescence intensity, *I* is the final fluorescence intensity, *a* is the saturation value at infinite time, *t* is the time, and *τ* is the characteristic time constant (Table S2). The analysis revealed a rapid increase in fluorescence intensity, with a time constant of *τ* = 32.4 sec, indicating the time required for the DNA1-SWCNTs fluorescence to reach 63.2% of its maximum intensity value.

To evaluate sensitivity, we measured the dose-dependent fluorescence response of DNA1-SWCNTs to increasing concentrations of L-DOPA, ranging from 0.001 μM to 5000 μM (Figure 1D). The normalized fluorescence intensity response data were fitted to the Hill equation:^[78]^

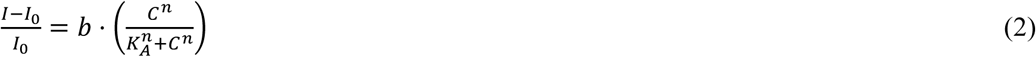

where *I_0_* is the initial fluorescence intensity, *I* is the final fluorescence intensity, *b* is the proportional coefficient, *C* is the analyte concentration, *n* is the Hill coefficient, and *KA* is the analyte concentration that results in 50% of the maximal response (Table S3). The LOD was calculated as three times the standard deviation of the SWCNT fluorescence intensity in the absence of L-DOPA, yielding an LOD of 0.003 μM. Previous studies have reported that the mean plasma concentration of L-DOPA in patients ranges from 2.28 μM to 35.85 μM, with peak levels reaching up to 69.72 μM.^[79]^ These concentrations exceed the LOD of DNA1-SWCNTs and remain within its dynamic range, indicating potential suitability for physiologically relevant conditions.

To enable future *in vivo* application of SWCNT-based sensors, a delivery matrix capable of encapsulating and transporting them is required. We therefore embedded the DNA1-SWCNTs within an injectable hydrogel that could be implanted in a minimally invasive manner.

FmocFF is a well-studied low molecular weight gelator known to self-assemble into fibrous networks, forming biocompatible hydrogels with tunable mechanical properties.^[80]^ We generated FmocFF hydrogels using the solvent-switch method, in which FmocFF dissolved in a water-miscible organic solvent, such as dimethyl sulfoxide (DMSO), is diluted with water, triggering molecular self-assembly into one-dimensional fibers that entangle to form a three-dimensional network, immobilizing the solvent, and thereby forming a hydrogel.^[81]^ However, when injected through a needle, these hydrogels failed to retain their hydrogel structure and underwent a liquid-solid phase separation. The applied shear stress during the injection process deformed the FmocFF network, and upon stress removal, the structure of the hydrogel did not recover. Instead, we observed solvent leakage, and when the injected hydrogel was oriented vertically at a 90° angle, it dripped, indicating inadequate mechanical resilience for injectable use (Figure 1E).

To improve the injectability of the hydrogel system, we explored the formation of composite hydrogels by combining FmocFF with additives,^[82–84]^ an approach previously shown to enhance structural and mechanical properties. We combined FmocFF with two biocompatible additives, the natural polysaccharides SA and HA, and evaluated their ability to reinforce the FmocFF network for enabling injectability. SA and HA were first dissolved in double-distilled water (ddw) followed by the addition of FmocFF dissolved in DMSO to initiate the gelation of the composite hydrogels via the solvent switch method. For each additive, we prepared hydrogel formulations with different FmocFF:additive mass ratios, namely, 80:20, 50:50, 20:80, while keeping the combined concentration of both the FmocFF and the additive constant at 5 mg mL^-1^. Upon injection, FmocFF/SA and FmocFF/HA hydrogels with the 80:20 and 50:50 ratios maintained their shape and hydrogel integrity, showing no leakage or displacement, even when tilted vertically (Figure 1F, G). In contrast, FmocFF/additive hydrogels with the 20:80 ratio lacked sufficient structural stability. Although they did not leak post-injection, they were easily displaced and failed to retain a defined shape when oriented vertically. Since neither HA nor SA forms hydrogels independently, we attribute this instability to the insufficient FmocFF fiber content at lower FmocFF ratios. Based on these results, we chose the 80:20 ratio FmocFF/additive hydrogels, which offer a stable and robust post-injection structural retention suitable for further development.

Having established both a responsive SWCNT suspension in solution and an injectable hydrogel formulation with favorable mechanical properties, we next integrated the DNA1-SWCNTs into the hydrogel matrix. To do so, the DNA1-SWCNTs suspension was mixed into either the SA or the HA solution prior to gelation. FmocFF was then added to initiate the self-assembly of the hydrogel. For the FmocFF:additive ratio of 80:20, the final hydrogel composition contained 4 mg mL^-1^ of FmocFF, 1 mg mL^-1^ of the additive (SA or HA), and 3 mg mL^-1^ of SWCNTs.

To evaluate the mechanical properties of the FmocFF/additive/SWCNT hydrogels, we conducted time sweep rheology measurements at a constant frequency of 1 Hz and a strain of 0.1%. These values were chosen based on frequency sweep and amplitude sweep tests, which confirmed that they lie within the linear viscosity region (LVR) (Figure S2). The storage modulus (G’) and the loss modulus (G’’) of the FmocFF/SA/SWCNT hydrogels increased over time, with G’ increasing in two steps before both moduli reached a plateau. For FmocFF/HA/SWCNT hydrogels, both moduli gradually increased until reaching a plateau (Figure 2A). In both cases, G’ exceeded G’’ throughout the measurement, indicating the formation of a stable hydrogel network.

**Figure 2.**
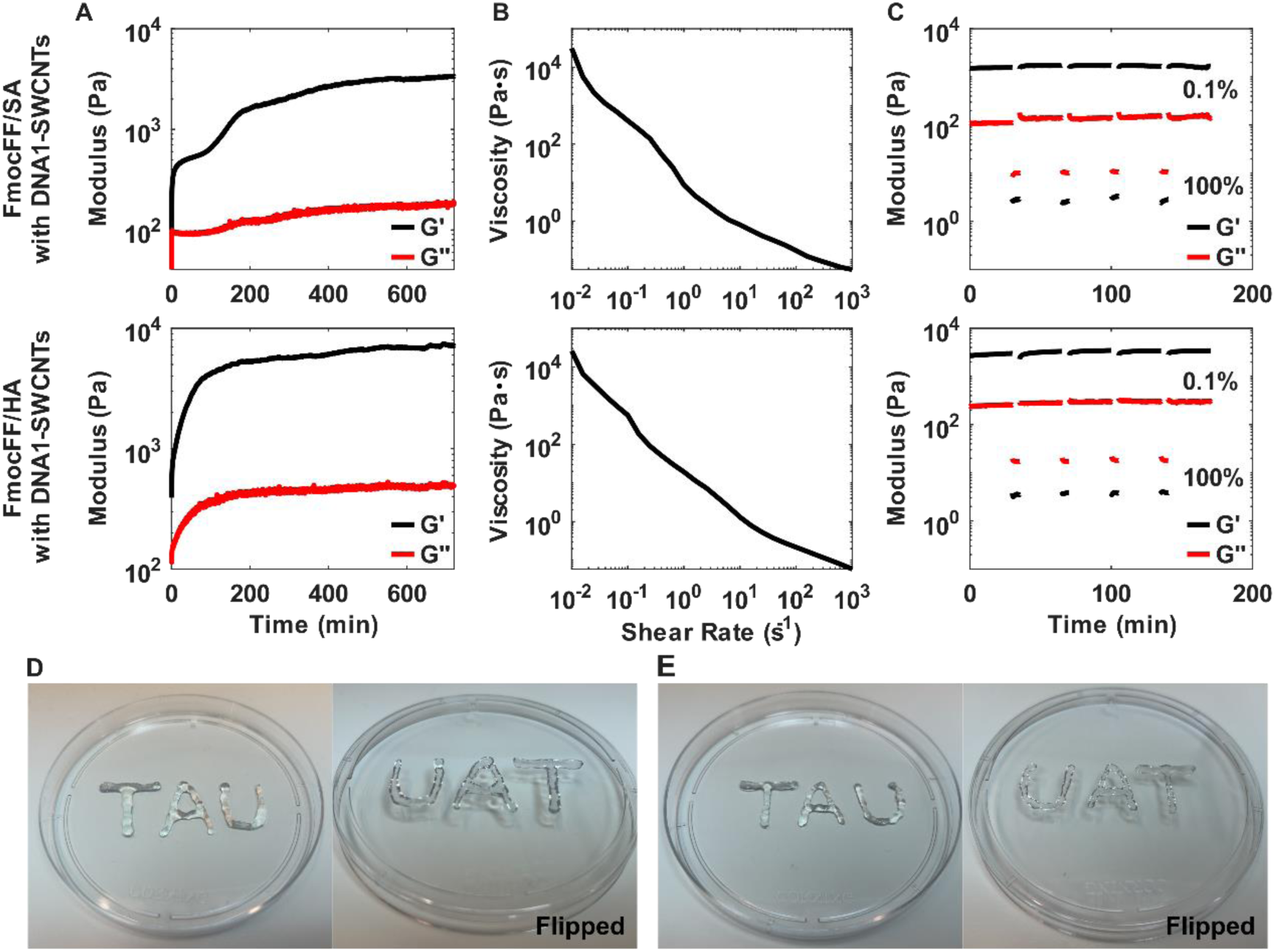
Rheological characterization of FmocFF/additive/SWCNTs hydrogels. (A) Time-sweep measurements. Storage modulus (G’, black line) and loss modulus (G’’, red line). (B) Flow-sweep measurements. (C) Thixotropic measurements alternating between 0.1% strain and 100% strain. Top row: FmocFF/SA/SWCNTs hydrogels. Bottom row: FmocFF/HA/SWCNTs hydrogels. (D) Photographs demonstrating the injectability of FmocFF/SA/SWCNTs hydrogels and (E) FmocFF/HA/SWCNTs hydrogels.

To function as an injectable carrier, a self-assembled hydrogel must exhibit both shear-thinning and self-healing abilities, enabling it to deform under shear stress (e.g., during injection), pass through a narrow needle, and subsequently recover its structural integrity upon the removal of shear stress.^[85]^ We first assessed the shear-thinning property by conducting flow sweep rheological measurements, during which we measured the viscosity of the hydrogels at increasing shear rates. The viscosity of both FmocFF/additive/SWCNT hydrogels decreased with increasing shear rates, confirming shear-thinning behavior (Figure 2B). We next evaluated the hydrogels self-healing capacity via thixotropic testing. The hydrogels were alternately subjected to a low 0.1% strain, coinciding with the LVR of the hydrogels, for 1800 seconds, and a high 100% strain for 300 seconds to disrupt the hydrogel structure (Figure 2C). During the low-strain phase, the storage modulus G’ exceeded the loss modulus G’’, indicating a gel state. However, upon application of the high strain, G’’ exceeded G’, signifying a transition to a sol state. When the low strain was reapplied, both G’ and G’’ returned to their initial values, demonstrating the hydrogel’s ability to self-recover. This reversible behavior was sustained over four cycles, indicating robust self-healing capacity under high strain, an essential feature for repeated mechanical stress encountered during injection. The practical injectability of the self-assembled hydrogels was further validated by injecting them through a syringe needle (Figure 2D, E). Post injection, the FmocFF/additive/SWCNT hydrogels maintained their shape and gel integrity without displacement, even when flipped, showcasing their suitability as injectable platforms for SWCNT sensor delivery.

To confirm the successful incorporation and spatial distribution of the SWCNTs within the hydrogel matrix, we employed NIR fluorescence microscopy to capture the fluorescence emission of the DNA1-SWCNTs and overlaid it on a bright field image of the hydrogels to visualize the SWCNT localization (Figure S3). In both injected FmocFF/SA/SWCNT and FmocFF/HA/SWCNT hydrogels, the fluorescent SWCNTs appeared to be well-dispersed and closely associated with the fibrous network, suggesting effective integration into the hydrogel architecture.

After incorporating DNA1-SWCNTs into FmocFF/additive hydrogels, we evaluated their sensitivity to L-DOPA by performing dose-dependent measurements. The FmocFF/additive/SWCNTs hydrogels were deposited in a 96-well plate and allowed to gel for 24 hours. An equal volume of FmocFF/additive hydrogel containing L-DOPA (FmocFF/additive/L-DOPA) was then layered on top of the preformed FmocFF/additive/SWCNTs hydrogel by injection (Figure 3A). In this configuration, the effective L-DOPA concentration across the system was half that of the injected FmocFF/additive/L-DOPA hydrogels, yielding a final concentration range of 0.001 μM to 5000 μM (Figure 3B). The normalized fluorescence intensity response data were fitted by the Hill equation (Figure 3C, Table S3). The calculated LOD for both additives was 0.0018 μM for the DNA1-SWCNTs embedded in the FmocFF/SA hydrogels and 0.3828 μM for the DNA1-SWCNTs embedded in the FmocFF/HA hydrogels. Notably, the reported mean plasma concentrations of L-DOPA in patients remain well above these LOD values and within the dynamic range of the embedded DNA1-SWCNTs in both hydrogel systems. These results confirm that DNA1-SWCNTs maintain their sensitivity to L-DOPA even when embedded within a hydrogel matrix, highlighting the robustness of this sensing platform.

**Figure 3.**
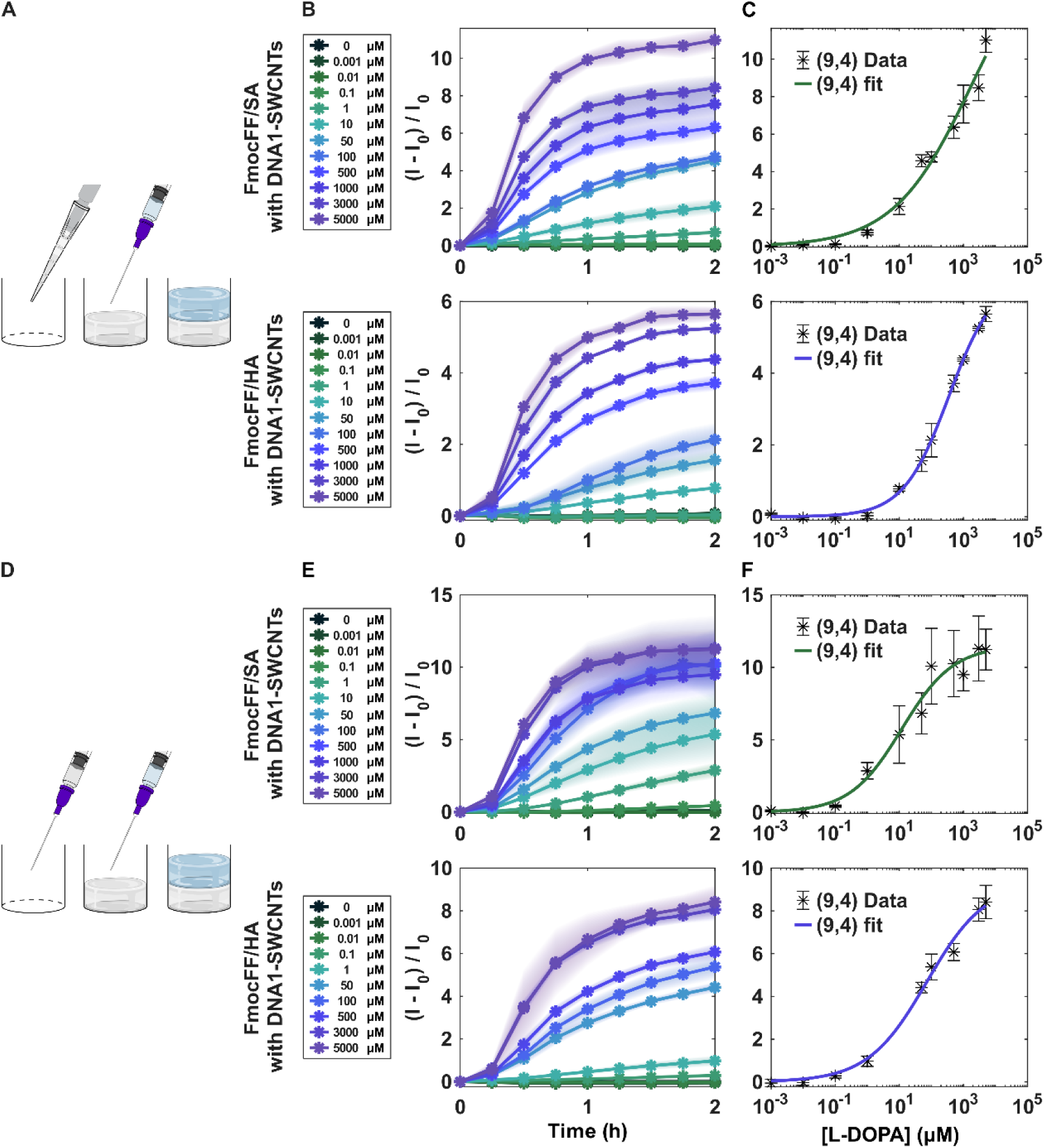
Fluorescence response of embedded DNA1-SWCNTs in FmocFF/additive hydrogels to L-DOPA. (A) Scheme of the experimental protocol. Gray represents FmocFF/additive hydrogels with DNA1-SWCNTs, and blue represents the FmocFF/additive with L-DOPA. FmocFF/additive/SWCNTs hydrogel solution was cast into a 96-well plate and left to gel overnight. FmocFF/additive/L-DOPA preformed hydrogel was then injected on top of the formed FmocFF/additive/SWCNTs hydrogel. (B) Time-dependent normalized fluorescence response of the DNA1-SWCNTs in the pipetted hydrogels. (C) Dose-dependent normalized fluorescence response of the (9,4) chirality of the DNA1-SWCNTs in the pipetted hydrogels to increasing concentrations of L-DOPA, fitted by the Hill equation (Equation 2, colored line).

Top row: FmocFF/SA hydrogels with DNA1-SWCNTs. Bottom row: FmocFF/HA hydrogels with DNA1-SWCNTs. (D) Scheme of the experimental protocol. Gray represents FmocFF/additive hydrogels with DNA1-SWCNTs, and blue represents the FmocFF/additive with L-DOPA. FmocFF/additive/SWCNTs preformed hydrogel was injected into a 96-well plate. FmocFF/additive/L-DOPA preformed hydrogel was then injected on top of the injected FmocFF/additive/SWCNTs hydrogel. (E) Time-dependent normalized fluorescence response of the DNA1-SWCNTs in the injected hydrogels. (F) Dose-dependent normalized fluorescence response of the (9,4) chirality of the DNA1-SWCNTs in the injected hydrogels to increasing concentrations of L-DOPA, fitted by the Hill equation (Equation 2, colored line). Top row: FmocFF/SA hydrogels with DNA1-SWCNTs. Bottom row: Injected FmocFF/HA hydrogels with DNA1-SWCNTs.

Having established that DNA1-SWCNTs embedded in hydrogels that were pipetted and then allowed to gel, retain a robust dose-dependent response to L-DOPA, we next examined whether this sensing capability is preserved in preformed hydrogels post-injection through a fine needle. The FmocFF/additive/SWCNTs hydrogels were loaded into syringes and left to gel inside for 24 hours. After the hydrogel was formed, it was injected into a 96-well plate. A layer of equal volume of FmocFF/additive/L-DOPA was injected on top of the injected preformed FmocFF/additive/SWCNTs hydrogels (Figure 3D), and the normalized fluorescence intensity response was measured for increasing concentrations of L-DOPA ranging from 0.001 μM to 5000 μM (Figure 3E). The data were fitted by the Hill equation (Figure 3F, Table S3), and the LOD was calculated to be 0.0071 μM for the DNA1-SWCNTs incorporated within the FmocFF/SA hydrogels and 0.1784 μM for the DNA1-SWCNTs in the FmocFF/HA hydrogels. These values remain well below reported plasma concentrations of L-DOPA in patients, indicating that the injection process has little to no effect on sensor performance. Thus, the encapsulated DNA1-SWCNTs preserve their sensitivity to L-DOPA even after hydrogel delivery, supporting the robustness of this injectable sensing platform.

The FmocFF/additive hydrogels provide an additional biocompatible interface for SWCNTs in a biological environment. To assess whether exposure to a complex biological fluid affects the fluorescence response of DNA1-SWCNTs to L-DOPA, we conducted dose-dependent experiments in hydrogels containing both L-DOPA and fetal bovine serum (FBS) (Figure 4A). The final L-DOPA concentration in the system ranged from 0.001 μM to 5000 μM (Figure 4B), and the normalized fluorescence intensity response data were fitted by the Hill equation (Table S3, Figure 4C). The calculated LOD was 0.5645 μM for the DNA1-SWCNTs encapsulated within the FmocFF/SA hydrogels and an LOD of 0.6167 μM for the DNA1-SWCNTs encapsulated within the FmocFF/HA hydrogels. Despite exposure to a complex biological environment, L-DOPA detection remained within physiologically relevant limits.

**Figure 4.**
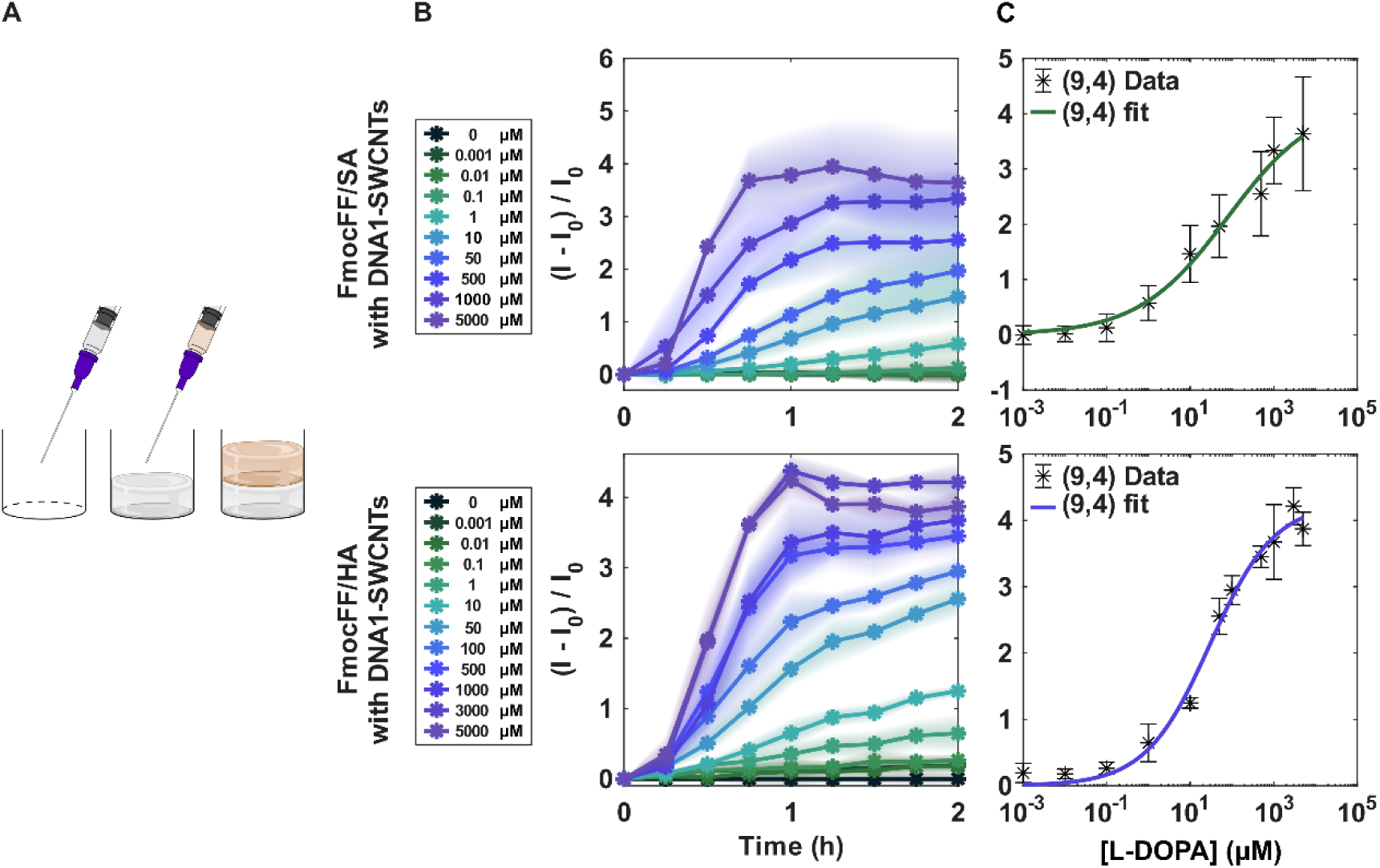
Fluorescence response of embedded DNA1-SWCNTs in injected FmocFF/additive hydrogels to L-DOPA with FBS. (A) Scheme of the experimental protocol. Gray represents the FmocFF/additive hydrogels with DNA1-SWCNTs, and brown represents the FmocFF/additive with L-DOPA and FBS. FmocFF/additive/SWCNTs preformed hydrogel was injected into a 96-well plate. FmocFF/additive with L-DOPA and FBS preformed hydrogel was then injected on top of the injected FmocFF/additive/SWCNTs hydrogel. (B) Time-dependent normalized fluorescence response of the DNA1-SWCNTs in the injected hydrogels. (C) Dose-dependent normalized fluorescence response of the (9,4) chirality of the DNA1-SWCNTs in the injected hydrogels to increasing concentrations of L-DOPA with FBS, fitted by the Hill equation (Equation 2, colored line). Top row: Injected FmocFF/SA hydrogels with DNA1-SWCNTs. Bottom row: Injected FmocFF/HA hydrogels with DNA1-SWCNTs.

To probe the performance of the FmocFF/additive/SWCNTs delivery platform in a relevant tissue-like setting, hydrogels containing DNA1-SWCNTs were injected into a tissue-mimicking phantom designed to reproduce the mechanical properties and optical scattering of subcutaneous tissue.^[86]^ The phantom hydrogel consisted of gelatin and agar to emulate the viscoelasticity of the subcutaneous layer, together with a lipid emulsion to mimic scattering from spherical lipid droplets, the dominant scatterers in subcutaneous tissue^[87]^. The gelatin–agar–lipid mixture was cast in a 24-well plate and allowed to gel. A small volume of the FmocFF/additive/SWCNTs hydrogel, which constitutes 3.33% of the volume of the phantom, was then injected into the phantom (Figure 5A). An L-DOPA solution with a volume equal to that of the phantom was subsequently deposited on top of the phantom, yielding final effective L-DOPA concentrations ranging from 70 μM to 5000 μM. Under these tissue-like conditions, a concentration-dependent increase in fluorescence intensity was observed for SWCNTs integrated into both of the FmocFF/additive hydrogels (Figure 5B).

**Figure 5.**
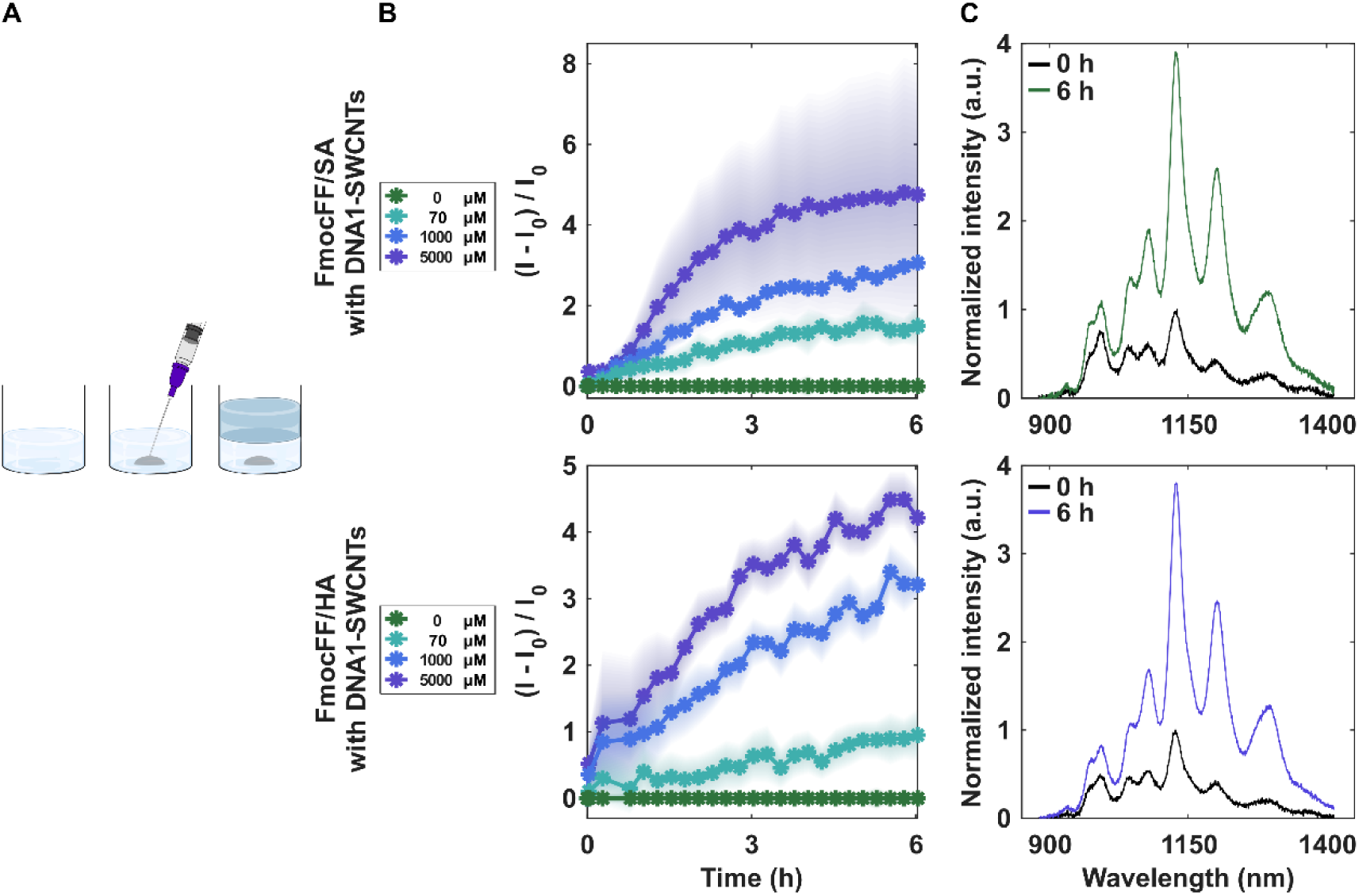
Fluorescence response of DNA1-SWCNTs in FmocFF/additive hydrogels to L-DOPA after injection into a subcutaneous mimicking phantom. (A) Scheme of the experimental protocol. Gray represents the FmocFF/additive hydrogels with DNA1-SWCNTs, light blue represents the tissue mimicking phantom, and blue represents the L-DOPA solution. The hydrogel phantom was cast into a 24-well plate and was left to gel overnight. FmocFF/additive/SWCNTs preformed hydrogel was injected into the hydrogel phantom. An L-DOPA solution was then deposited on top of the injected hydrogel phantom. (B) Time-dependent normalized fluorescence response of the DNA1-SWCNTs in FmocFF/SA hydrogels injected into a phantom. (C) Normalized intensity of DNA1-SWCNTs in FmocFF/SA hydrogels injected into a phantom before (black line) and after (colored line) the addition of L-DOPA to the phantom.

The scattering elements can, in principle, modify the optical path. However, scattering in the subcutaneous layer is more significant in the visible light (400 nm – 700 nm) compared to the NIR range (700 nm – 1500 nm),^[87]^ so its influence on the fluorescence emitted from the SWCNTs (900 nm – 1400 nm) should be less notable. Consistently, and despite the scatterers in the phantom, we still observed a clear fluorescent increase at 70 μM of L-DOPA (Figure 5C), a physiologically relevant plasma concentration in patients. These results demonstrate that the sensitivity of the embedded DNA1-SWCNTs is preserved even after hydrogel injection into an environment that simulates a biological tissue, highlighting the potential of this platform for *in situ* sensing applications.

## Conclusion

In this study, we introduced a minimally invasive platform for delivering SWCNT-based NIR fluorescent sensors using injectable, self-assembling, preformed peptide hydrogels. By incorporating DNA-suspended SWCNTs into FmocFF-based hydrogels supplemented with SA or HA, we formed a three-dimensional network of shear-thinning, self-healing materials that can be injected through a fine needle while retaining their structural integrity and sensor functionality. These composite hydrogels enable analyte diffusion throughout the network, allowing the encapsulated SWCNTs to interact with the target molecule, which in turn, induces a robust fluorescence response. Using L-DOPA, the primary treatment for Parkinson’s disease, as a model analyte, we demonstrated a sensitive, dose-dependent response within physiologically relevant plasma concentrations. Importantly, this sensing capability was preserved after hydrogel injection, in the presence of serum, and within a subcutaneous tissue-mimicking phantom that reproduces both the mechanical and scattering properties of soft tissue, underscoring the robustness and adaptability of the system under tissue-like optical conditions.

This injectable hydrogel platform offers a versatile route for the localized deployment of SWCNT-based sensors, enabling non-surgical, *in situ* monitoring of drug concentrations. Beyond L-DOPA, the approach can be extended to a wide array of SWCNT-based sensors for other drugs and clinically relevant biomarkers, opening opportunities for personalized therapeutic drug monitoring. The strategy highlights how self-assembling peptide hydrogels can serve as biocompatible carriers for nanosensors, advancing continuous monitoring technologies that could aid clinicians in tailoring treatments, tracking disease progression, and ultimately improving patient care.

## Experimental Section

### Materials

Levodopa (L-DOPA), hyaluronic acid (HA), sodium alginate (SA), and gelatin (type A, 300 grams bloom) were purchased from Sigma-Aldrich. Agar was purchased from Becton Dickinson. Fluorenylmethoxycarbonyl (Fmoc)-diphenylalanine (FmocFF) was purchased from Bachem. HiPCO single-walled carbon nanotubes (SWCNTs) were purchased from NanoIntegris. Single-stranded DNA (ssDNA) oligonucleotides (Table S1) were purchased from Integrated DNA Technologies. Fetal bovine serum (FBS) was purchased from Sartorius. Intralipid-20% was purchased from MedChemExpress.

### Single-walled carbon nanotube suspension

SWCNTs (1 mg) were suspended with ssDNA (2 mg) in an NaCl solution (0.1 M). The mixture was first bath sonicated (Elma P-30H, 80 Hz, 10 min) and then tip sonicated in an ice bath (QSonica, Q125, 3mm tip, 4 W, 40 min). The resulting suspension was centrifuged twice (16,100 rcf, 90 min). After each centrifugation, 80% of the supernatant was collected, and the pellet was discarded. The suspension was then transferred to 100 kDa ultra centrifugal filters (Merck Millipore) and centrifuged (14,000 rcf, 30 min) to remove excess ssDNA oligonucleotides. The ssDNA-SWCNTs were redispersed in double-distilled water (ddw) and centrifuged again (16,100 rcf, 30 min), after which 80% of the supernatant was collected and the pellet discarded. The absorption spectra of the ssDNA-SWCNT suspensions were measured using a UV-Vis-NIR spectrophotometer (Shimadzu UV-3600 Plus) in the wavelength range of 200 nm – 1400 nm. The concentration of the suspended ssDNA-SWCNT was determined by the absorption value at 632 nm using an extinction coefficient of ε_632 nm_ = 0.036 L mg^-1^ cm^-1^.^[74]^

### Hydrogel formation

A stock solution of HA and SA in ddw (5 mg mL^-1^) was made by rotating the mixtures overnight until complete dissolution. The FmocFF/additive hydrogels were formed *via* the solvent-switch method, as described below. First, the additive stock solution was added to ddw and gently mixed to reach the final desired concentration of the additive in the hydrogel. FmocFF, dissolved in dimethyl sulfoxide (DMSO, 100 mg mL^-1^), was then added to the diluted additive solution at defined FmocFF:additive ratios of 20:80, 50:50, 80:20, or 100:0, with a combined concentration of 5 mg mL^-1^. Upon FmocFF addition, the mixture was vortexed and left undisturbed to gel. Specifically, the FmocFF/additive hydrogels with an 80:20 ratio, used as the primary formulation in this work, contained 4 mg mL^-1^ of FmocFF and 1 mg mL^-1^ of the additive.

FmocFF/additive/SWCNT hydrogels and FmocFF/additive/L-DOPA hydrogels were prepared similarly to the FmocFF/additive hydrogels described above. The additive stock solution was first added to a solution of either ssDNA-SWCNT or L-DOPA, dissolved in ddw, and gently mixed. For FmocFF/additive/L-DOPA hydrogels with 10% FBS, FBS was added to the additive and L-DOPA solution, and the mixture was gently mixed. FmocFF, dissolved in DMSO (100 mg mL^-1^), was then added to initiate the gelation. The final concentration of ssDNA-SWCNT in the hydrogels was 3 mg mL^-1^, and the final concentration of L-DOPA in the hydrogels ranged from 0.002 μM to 10,000 μM. When FBS was added, its volume was 10% of the final hydrogel volume. In all formulations, the combined concentration of FmocFF and the additive was maintained at 5 mg mL^-1^.

### Subcutaneous tissue mimicking phantom formation

The tissue-mimicking phantom was prepared as previously described.^[86]^ Agar powder was added to water and heated to 95°C until completely dissolved. Gelatin powder was added to cold water and left to rest for 1 hour. The gelatin mixture was then heated to 37°C until completely dissolved. The agar and gelatin solutions were combined with an Intralipid 20% emulsion to a final concentration of 0.2% w/v of agar, 2% w/v of gelatin, and 7.8% v/v of intralipid-20%. The mixed solution was deposited into 24-well plates and left to gel overnight at 4°C.

### Fluorescence NIR spectroscopy

Fluorescence emission spectra were recorded using a 96-well plate mounted on the stage of an inverted microscope (Olympus IX73). A super-continuum white-light laser (NKT-photonics, Super-K Extreme) with a bandwidth filter (NKT-photonics, Super-K Varia, Δλ = 20 nm) was used as the excitation light source. Spectra were acquired at an excitation wavelength of 730 nm with an excitation power of 20 mW. Fluorescence emission was spectrally resolved using a spectrograph (Spectra Pro HRS-300, Princeton Instruments) with a 500 μm slit-width and a 150 g mm^-1^ grating, and recorded with an InGaAs camera (PylonIR, Teledyne Princeton Instruments). Excitation-emission maps were generated using excitation wavelengths ranging from 500 nm to 800 nm in 2 nm increments. Fluorescence intensity changes were quantified by fitting the chirality peaks with a Lorentzian function and adjusted to 0 mM concentration. The intensity modulations in dose-dependent experiments were calculated after a two-hour incubation.

### Fluorescence microscopy

Images were acquired using an inverted fluorescence microscope (Olympus IX83). Bright-field transmission images were obtained under an LED light source illumination and captured with an EMCCD camera (Andor, iXon Ultra). SWCNT fluorescence was excited using a 730 nm continuous-wave laser (MDL-MD-730 1.5W, Changchun New Industries) at an excitation power of 500 mW. The laser excitation light was directed onto the sample using a dichroic mirror (900 nm lp, chroma, T900lpxxrxt), and the NIR emission of the SWCNTs was detected with an InGaAs camera (Raptor, Ninox 640 VIS-SWIR) after passing through an additional 900 nm long-pass emission filter (Chroma, ET900lp). A UPLFN 100X objective, 1.3 NA was used. All images were processed by Fiji (ImageJ) and MATLAB.

### Rheometry

Rheology measurements of the FmocFF/additive hydrogels with SWCNTs were conducted using a rotational rheometer (Discovery HR2, TA Instruments, USA) equipped with a 20 mm parallel plate geometry. Time-sweep oscillatory measurements were carried out for 12 h at 0.1% strain and a frequency of 1 Hz. These conditions were confirmed to be in the linear viscoelastic region of the hydrogels. Frequency sweep oscillatory measurements were conducted at a 0.1% strain over a frequency range of 0.01 Hz – 100 Hz. Amplitude sweep oscillatory measurements were performed at a strain range of 0.01% – 100% with a frequency of 1 Hz. Flow sweep measurements were carried out at a shear rate of 0.01 s^-1^ – 1000 s^-1^. Thixotropic tests were conducted at a frequency of 1 Hz, alternating between 30 minutes at 0.1% strain and 5 minutes at 100% strain. All measurements were performed at 25°C.

## Acknowledgements

G.B. acknowledges the support of the Zuckerman STEM Leadership Program, the ERC NanoNonEq (Grant No. 101039127), the Israel Science Foundation (Grant no. 196/22), the Ministry of Science, Technology, and Space, Israel (Grant nos. 1001818370 and 0008452), the Marian Gertner Institute for Medical Nanosystems at Tel Aviv University, and the Naomi Prawer Kadar Foundation. S.K. thanks the Aufzien Family Center for the Prevention and Treatment of Parkinson’s Disease at Tel Aviv University for the Prof. Nir Giladi Legacy Scholarship, as well as the Cancer Biology Research Center (CBRC) at Tel Aviv University for the Esther Weinstat Graduate Student Award.

---

Injectable peptide hydrogels offer a minimally invasive delivery platform for single-walled carbon nanotube (SWCNT)-based near-infrared fluorescent sensors, enabling real-time measurement of drug concentrations. Encapsulated SWCNTs show a dose-dependent fluorescence response across physiologically relevant concentrations, and retain their fluorescence response post-injection, in the presence of serum, and within a tissue phantom.

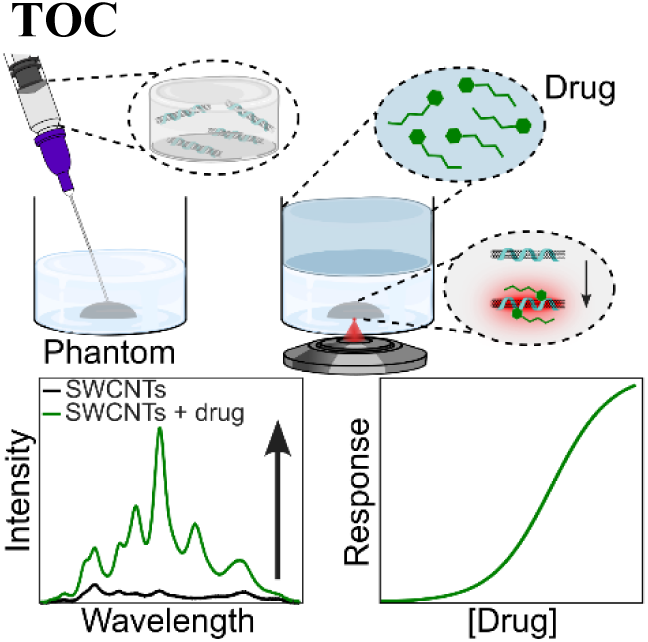

