## Supporting Information for "Minimally invasive delivery of optical nanosensors using injectable hydrogels"

Table S1. ssDNA sequences used for suspending SWCNTs.

| **Name** | **Sequence** |
|---|---|
| $(AT)_{15}$ | ATATATATATATATATATATATATATATAT |
| $(ATTT)_{7}$ | ATTTATTTATTTATTTATTTATTTATTT |
| $C_{30}$ | CCCCCCCCCCCCCCCCCCCCCCCCCCCCCC |
| $(GT)_{15}$ | GTGTGTGTGTGTGTGTGTGTGTGTGTGTGT |
| $(GC)_{30}$ | GCGCGCGCGCGCGCGCGCGCGCGCGCGCGCG<br>CGCGCGCGCGCGCGCGCGCGCGCGCGCGC |
| DNA1:<br>$(GT)_{15}$GATCTAAGGCGTGTAT | GTGTGTGTGTGTGTGTGTGTGTGTGTGTGTGA<br>TCTAAGGCGTGTAT |
| DNA2:<br>$(GT)_{15}$GATGCGTGTCTAAGAT | GTGTGTGTGTGTGTGTGTGTGTGTGTGTGTGA<br>TGCGTGTCTAAGAT |

Table S2. Fit parameters and 95% confidence intervals of data fitted to equation 1.

| | **a** | **τ (sec)** | **R-square** |
|---|---|---|---|
| Figure 1C | 18.3865 (18.2054, 18.5676) | 32.4 (22.32, 42.84) | 0.9987 |

Table S3. Fit parameters and 95% confidence intervals of data fitted to equation 2.

| | **b** | **$K_A$ (μM)** | **n** | **R-square** |
|---|---|---|---|---|
| Figure 1D | 27.46 (15.54, 39.39) | 213.2 (-426.7, 853) | 0.3371 (0.1862, 0.4881) | 0.9843 |
| Figure 3C, top | 17.33 (1.318, 33.33) | 1858 (-8035, 11750) | 0.3551 (0.1544, 0.557) | 0.9813 |
| Figure 3C, bottom | 6.734 (6.136, 7.333) | 356.2 (220, 492.5) | 0.6081 (0.5314, 0.6847) | 0.9989 |
| Figure 3F, top | 11.49 (9.776, 13.2) | 11.9 (-0.145, 23.95) | 0.539 (0.2896, 0.7884) | 0.9786 |
| Figure 3F, bottom | 9.37 (7.7, 11.04) | 72.43 (-1.616, 146.5) | 0.471 (0.302, 0.6401) | 0.9938 |
| Figure 4C, top | 4.27 (2.802, 5.738) | 80.06 (-98.98, 259.1) | 0.4052 (0.2111, 0.5993) | 0.9865 |
| Figure 4C, bottom | 4.263 (3.766, 4.773) | 29.87 (9.611, 50.12) | 0.5615 (0.3782, 0.7448) | 0.9902 |

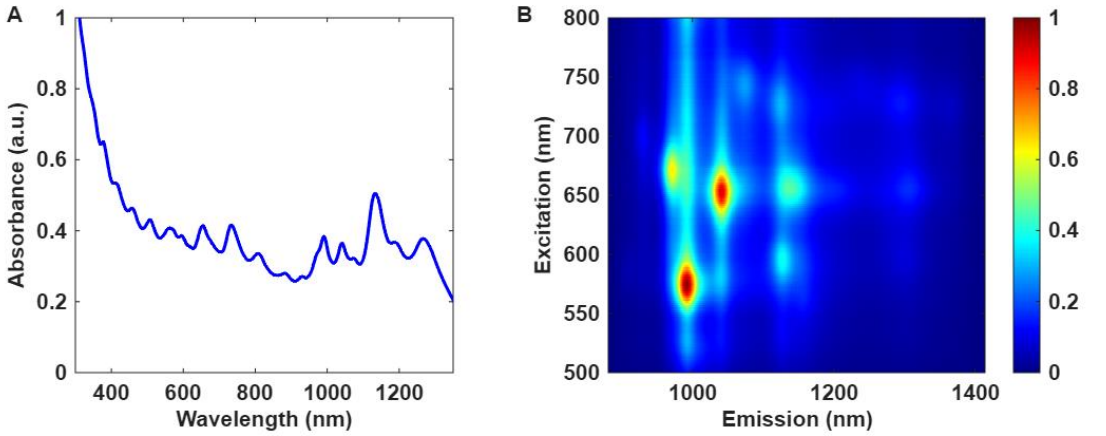


Figure S1. Characterization of DNA1-SWCNTs. (A) Absorption spectrum and (B) Excitation-emission map.

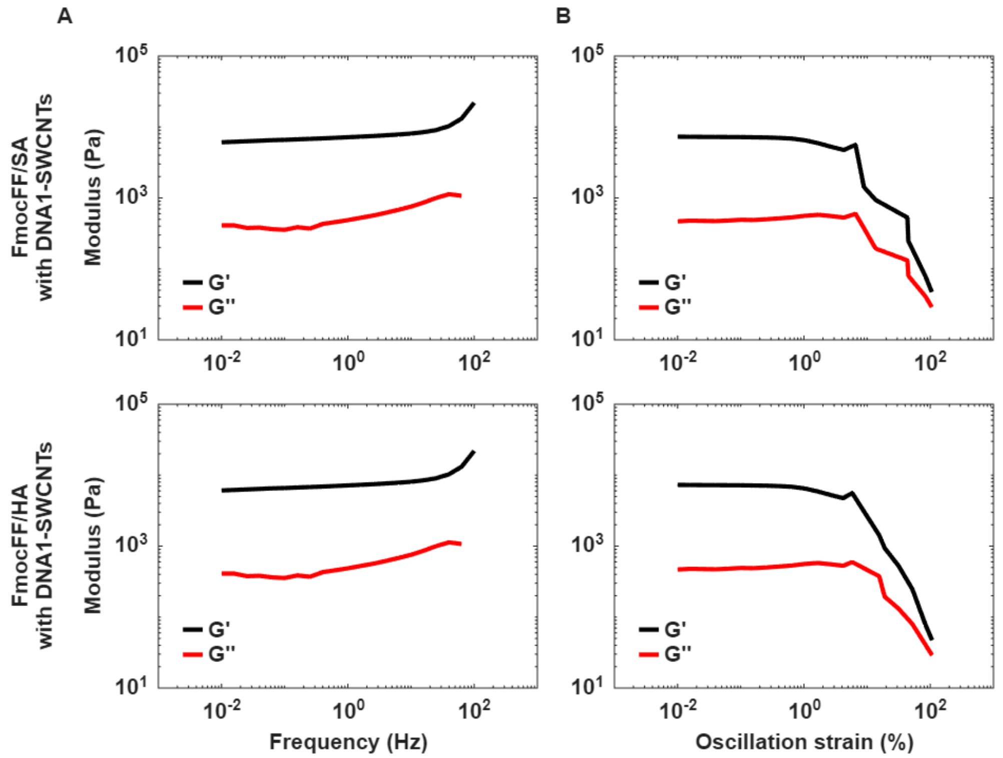


Figure S2. Rheological measurements of FmocFF/additive/SWCNTs hydrogels. (A) Frequency sweep rheology measurements. (B) Amplitude sweep rheology measurements. Storage modulus (G’, black line) and loss modulus (G’’, red line). Top row: FmocFF/SA/SWCNTs hydrogels. Bottom row: FmocFF/HA/SWCNTs hydrogels.

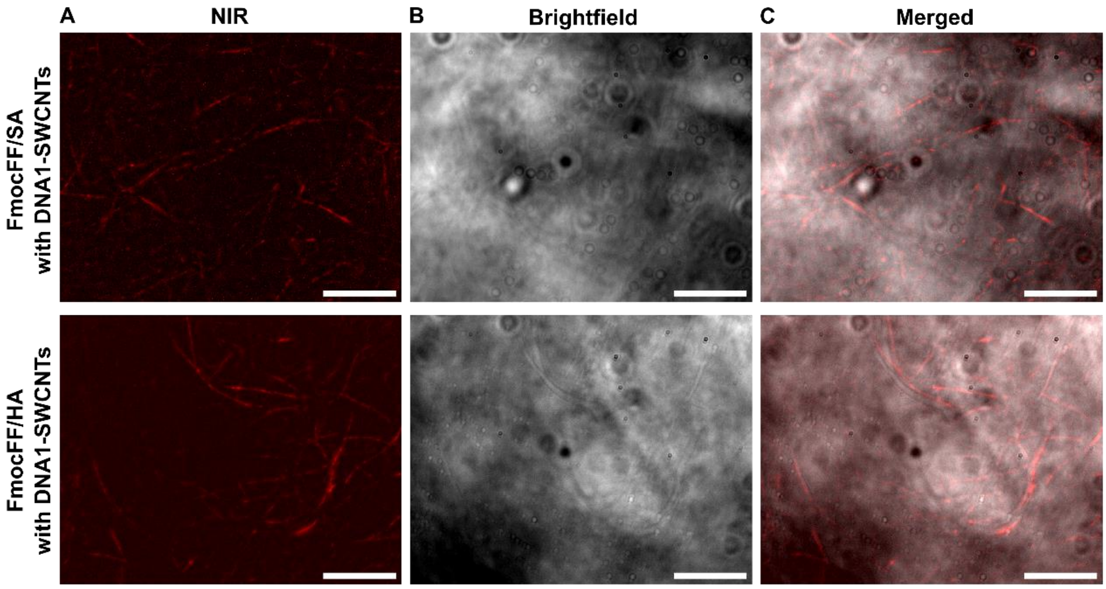


Figure S3. NIR fluorescence and brightfield imaging of FmocFF/additive/SWCNTs hydrogels. (A) NIR fluorescence imaging of DNA1-SWCNT integrated within the hydrogels. (B) Brightfield images of the FmocFF/additive/SWCNTs hydrogels. (C) Overlay of the NIR SWCNT fluorescence on top of the brightfield image. Top row: FmocFF/SA/SWCNTs. Bottom row: FmocFF/HA/SWCNTs. Scale bar: 20 µm.
